# Strain-specific galactose utilization by commensal *E. coli* mitigates *Salmonella* establishment in the gut

**DOI:** 10.1101/2025.03.03.641131

**Authors:** Christopher Schubert, Jana Näf, Lisa Petukhov, Leanid Laganenka, Yassine Cherrak, Wolf-Dietrich Hardt

**Author notes:** Correspondence (Y.C), (W.-D.H). Co-last authors (Yassine Cherrak, Wolf-Dietrich Hardt).

## Abstract

*Salmonella enterica* serovar Typhimurium (*S*. Tm) is a major cause of gastrointestinal diseases worldwide. To date, options for prevention or curative therapy remain limited. The gut microbiota plays a protective role against enteric diseases, particularly in preventing establishment and proliferation of *S*. Tm. While most research has focused on microbiota-mediated pathogen exclusion during the later, inflammation-dominated stages of infection, little is known about how microbiota members mitigate *S*. Tm early gut colonization. To address this gap, we conducted 24 h *in vivo* competitive experiments using *S*. Tm and different commensal *E. coli* strains. We observed a significant reduction in pathogen load, which was strain-specific and particularly evident with *E. coli* 8178. To investigate the underlying molecular mechanisms, we performed an *in vivo* screen using a rationally designed *S*. Tm library -which includes a wide range of carbohydrate utilization mutants - both in the absence and presence of *E. coli* strains. Our findings revealed that *E. coli* 8178-mediated *S*. Tm competition was driven by the exploitation of galactose during the early stage of infection. Identifying galactose as a key metabolite in pathogen exclusion by gut microbiota members enhances our mechanistic understanding of microbiota-mediated protection and opens new avenues for developing microbiota- and dietary-based strategies to better control intestinal infections.

## Introduction

Foodborne diarrhoea is a widespread disease, affecting 1 in 10 people globally and leading to over 420,000 deaths annually (WHO). Bacterial enteric pathogens are among the leading causes of foodborne diseases, with *Salmonella* responsible for 180 million cases of gastroenteritis each year (Besser, 2018). *Salmonella enterica* serovar Typhimurium (*S*. Tm) is among the most common non-typhoidal serovars that affect humans (Herzog et al., 2023, European Food Safety Authority). Research using antibiotic-pretreated specific pathogen free (SPF) C57BL/6J and gnotobiotic mouse models have identified two distinct growth phases for *S*. Tm in the gut (Wotzka et al., 2017). In these models, *S*. Tm establishes and grow in the intestinal lumen through monosaccharide-fuelled mixed acid fermentation during the early stage of infection (0 to 48 hours post-infection) (Nguyen et al., 2020; Nguyen et al., 2024; Schubert et al., 2025; Rogers et al., 2024). Subsequently, *S*. Tm employs the type III secretion systems encoded within the pathogenicity island 1 (T3SS-1) and 2 (T3SS-2) to trigger inflammation, creating conditions that favour proliferation, evolution and transmission of the pathogen (Lupp et al., 2007; Stecher et al., 2007; Winter et al., 2010; Thiennimitr et al., 2011; Stecher et al., 2012; Maier et al., 2013; Faber et al., 2017; Gillis et al., 2018; Nguyen et al., 2020; Fattinger et al., 2021; Rogers et al., 2021; Nguyen et al., 2024).

Under homeostatic conditions, the gut microbiota plays a variety of functions, including restricting the invasion of enteric pathogens (Bohnhoff & Miller, 1962). This phenomenon is referred to as colonization resistance and can often be attributed to specific commensal stains exhibiting pathogen-displacing capabilities (Buffie & Pamer 2013; Caballero-Flores et al., 2019). This is particularly relevant for *S*. Tm infections, where several microbiota members were shown to act as protective strains (Brugiroux et al., 2016; Sorbara & Pamer, 2019; Rogers et al., 2021; Spragge et al., 2023). The prevention of pathogen growth by intestinal commensals can occur through interference mechanisms involving the production of toxic compounds. This is exemplified by commensal *E. coli* or *Bacteroides* spp. which produce toxins and metabolites directly impacting *Salmonella* growth in the gastrointestinal tract (Sassonne-Corsi et al., 2016; Jacobson et al., 2018; Cherrak et al., 2024a). Alternatively, commensal bacteria can engage in competitive interactions by exploiting available nutrients, thereby reducing the fitness of their competitors – a mechanism known as nutrient exploitation. This holds true for the human probiotic *E. coli* Nissle strain, which restricts intestinal *S*. Tm growth by scavenging iron (Deriu et al., 2013). Besides iron, several electron acceptors such as oxygen, fumarate and nitrate were also shown to be actively used by commensal *E. coli* strains, thereby limiting *Salmonella*’s fitness in the gut (Litvak et al., 2019; Velazquez et al., 2019, Nguyen et al., 2020, Schubert et al, 2021, Liou et al., 2022). While the mechanisms driving *S*. Tm restriction by commensals are increasingly understood, most of this research focuses on the later stages of infection, particularly when inflammation is already established. In contrast, our understanding of how commensals restrict pathogen during the initial growth phase remains scarce, with galactitol standing as one of the few known carbohydrates exploited by commensal *E. coli* to limit *S*. Tm gut invasion (Eberl *et al*., 2021)

In this study, we aimed to address these gaps by investigating how commensal *Enterobacteriaceae* restrict *S*. Tm during the early phase of intestinal growth and identifying the carbohydrates driving this exclusion relationship.

## Results

### *E. coli* 8178 limits early *S*. Tm gut establishment independently of interference mechanisms

*E. coli* 8178 is a commensal strain, originally isolated from the murine gut and capable of outcompeting *S*. Tm at the later stages of infection once inflammation is established (Stecher et al., 2012). This competitive advantage, visible in the severely inflamed gut between 72- and 96-hours post-infection (p.i) is attributed to an interference mechanism involving the synthesis and secretion of siderophore-bound toxins (Cherrak et al., 2024a; Cherrak et al., 2025). While investigating this phenomenon, we found out that *E. coli* 8178 was also able to restrict the growth of *S*. Tm at the early stage of infection (24 h p.i) when administered either prior to or concomitantly with *S*. Tm (Nguyen et al., 2020; Cherrak et al., 2024a). To confirm this observation, we inoculated streptomycin-pretreated 129S6/SvEvTac mice harbouring a specific pathogen-free (SPF) microbiota with a 1:1 mixture of *E. coli* 8178 and *S*. Tm SL1344. Mice co-inoculated with *E. coli* 8178 displayed a 50-fold reduction in faecal pathogen loads at 24 hours post-infection compared to mice infected with *S*. Tm alone (Figure 1A, B). This significant reduction during the initial stage of *S*. Tm infection, which remains uncharacterized, was the focus of our investigation.

**Figure 1:**
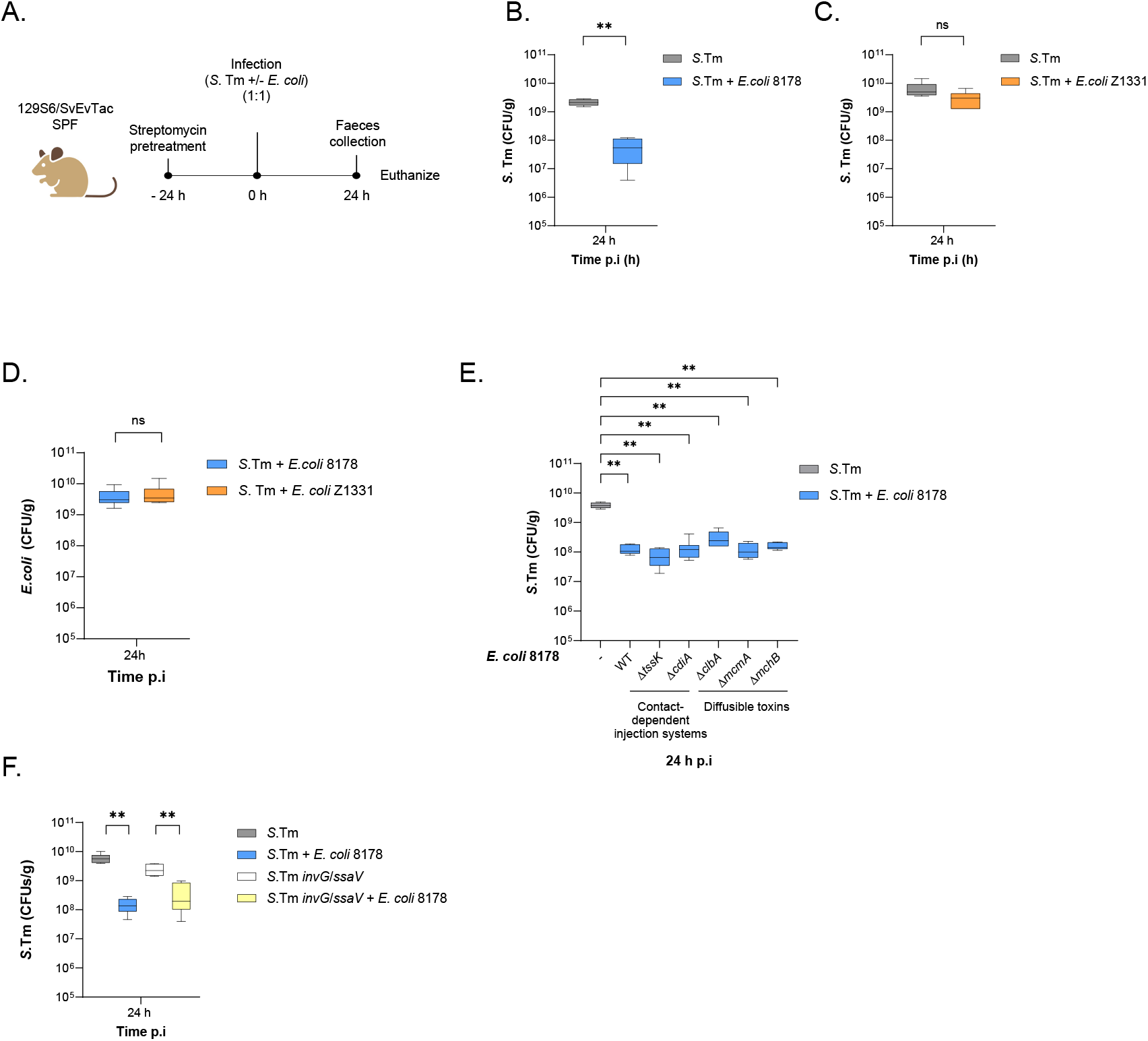
*E. coli* 8178*-*mediated *S*. Tm early growth restriction is independent of interfering competition interactions. **(A)** Experimental scheme. Streptomycin-pretreated specific pathogen-free (SPF) 129S6/SvEvTac mice were infected with either *S*. Tm alone or a 1:1 mixture of *S*. Tm and the indicated *E. coli* strain. The *E. coli* and *S*. Tm loads were determined by selective plating from faecal samples collected 24 h post-infection (p.i.) prior to euthanasia. Created in BioRender. Cherrak, Y. (2025) https://BioRender.com/n43x499. (**B-C**) *E. coli* mitigates the early growth of *S*. Tm in a strain-dependent manner. The *S*. Tm loads at 24 h p.i. are plotted and compared between *S*. Tm mono-infected and *S*. Tm + *E. coli* co-infected mice. *E. coli* 8178 (**B**) and Z1331 (**C**) were tested as competitors. (**D**) *E. coli* loads at 24 h p.i from Figures 1B and C are shown. (**E**) *E. coli* 8178 restricts *S*. Tm establishment independently of interfering mechanisms. The faecal load of *S*. Tm collected 24 h p.i. is plotted and compared between *S*. Tm mono-infected and *S*. Tm + *E. coli* 8178 co-infected mice. The *E. coli* 8178 mutant strains tested are listed and categorized based on the nature of the interference system disrupted. (**F**) Inflammation does not trigger *E. coli* 8178-mediated *S*. Tm competition 24 h p.i. *E. coli* 8178 competitiveness was tested against an avirulent *S*. Tm mutant (Δ*invG*/Δ*ssaV*). (**B-F**) All competitive experiments are presented in a box-and-whiskers plot, showing the minimum to maximum values, median, and interquartile range (25th to 75th percentiles). The bar plots show the median. Two-tailed Mann–Whitney U tests to compare 2 groups in each panel. *p* ≥ 0.05 not significant (ns), *p* < 0.05 (*), *p* < 0.01 (**). A minimum of 5 mice (n ≥ 5) were used for each experimental group. CFU: colony forming units.

First, we aimed to determine whether *S*. Tm growth attenuation at 24 hours p.i was specific to *E. coli* 8178 or if it was a common effect among other commensal *E. coli* strains. To explore this, we co-inoculated streptomycin-pretreated 129S6/SvEvTac mice with a mixture of *S*. Tm and a different *E. coli* strain. We selected *E. coli* Z1331, a recent and genetically tractable isolate from a healthy human volunteer (Wotzka et al., 2018). This strain has previously been used to study autoinducer-2-mediated chemotaxis in both interspecies and intraspecies signalling. (Laganenka et al., 2023, 2024). Interestingly, in this context, the competitive elimination of *S*. Tm at 24 hours p.i was not observed (Figure 1C, Supplementary Figure 1A) despite *E. coli* Z1331 shedding at similar levels as *E. coli* 8178 at 24 h p.i (Figure 1D). This indicated that *S*. Tm growth attenuation by *E. coli* 8178 is a strain-specific feature. Given the variety of antimicrobial systems encoded by *E. coli* 8178, we next hypothesized that *S*. Tm competition at 24 hours was driven by an interference mechanism, similar to what occurs during the late-stage infection, when inflammation peaks (Cherrak et al., 2024a; Cherrak et al., 2025). To test this, we systematically mutated each known interference factor in the *E. coli* 8178 genome, including both diffusible and contact-dependent toxin delivery machineries. We then assessed the ability of each *E. coli* 8178 mutant to displace *S*. Tm using streptomycin-pretreated SPF 129S6/SvEvTac mice and found that all strains remain effective in reducing *S*. Tm early gut establishment (Figure 1E). Finally, considering the variety of antagonistic relationships triggered during inflammation (Stecher et al., 2012; Deriu et al., 2013; Sassone-Corsi et al., 2016; Velazquez et al., 2019; Cherrak et al., 2024a), we examined whether *E. coli* 8178-mediated exclusion of *S*. Tm was dependent on an inflammatory response. To investigate this, we infected streptomycin-pretreated SPF 129S6/SvEvTac mice with *E. coli* 8178 and an isogenic *S*. Tm strain lacking both type three secretion systems (T3SS-1, Δ*invG* and T3SS-2 Δ*ssaV*), thus unable to trigger a severe gut inflammation (Stecher et al., 2007). Notably, the faecal loads of the *S*. Tm Δ*invG* Δ*ssaV* at 24 h p.i were consistently reduced by the presence of *E. coli* 8178 (Figure 1F). Overall, we concluded that *E. coli* 8178-mediated inhibition of *S*. Tm during the initial growth phase was independent of inflammation and did not rely on any known interference mechanisms.

### Impact of *E. coli* 8178 on *S*. Tm sugar metabolism during initial growth phase

Since exclusion of *S*. Tm by *E. coli* 8178 at 24 h post-infection is likely not due to an interference mechanism, we hypothesized that this effect may be explained by nutrient exploitation. Recently, hexoses have been identified as the primary nutrient source driving *S*. Tm colonization during the initial growth phase (Rogers et al., 2024; Nguyen et al., 2024; Schubert et al., 2025), making them a potential prime target for nutrient exploitation. To explore this further, we employed a targeted library of 35 *S*. Tm mutants, each defective in specific carbohydrate-utilizing enzymes (Schubert et al., 2025) and uniquely labelled with a chromosomal, fitness-neutral DNA barcode (WISH tag) (Daniel et al., 2024) (Figures 2A, B). We screened for metabolic-deficient *S*. Tm strains and tested whether mutant fitness was impacted by the presence of *E. coli* 8178. We also included *E. coli* Z1331 for comparison to highlight specific *E. coli* 8178-driven changes (Figure 2B). To control the quality of the generated pool data, seven wild-type (WT) strains carrying distinct WISH tags were included. This allowed to assess the bottleneck severity by calculating the Shannon evenness score, with mice scoring below 0.9 being excluded from the competitive index analysis (Maier et al., 2014; Schubert et al., 2025). Furthermore, two control *S*. Tm mutants (Δ*dcuABC*, Δ*frd*), which are deficient in fumarate respiration and exhibit an initial growth defect, were included as internal standards to validate our findings (Nguyen et al., 2020; Maier et al., 2013; Schubert et al., 2025). Streptomycin-pretreated 129S6/SvEvTac mice were infected with the targeted *S*. Tm carbohydrate-mutant pool, both in the absence and presence of *E. coli* 8178 or *E. coli* Z1331. The raw fitness data for each *S*. Tm metabolic mutants across these 3 conditions are presented in the Source data file. The Shannon evenness score and the fitness of the *S*. Tm control mutants for each dataset are presented in Supplementary Figures 2A and B. As previously reported, mutants deficient in fumarate respiration are already attenuated by day 1 post-infection (Schubert et al., 2025), further validating the reliability of our downstream analysis.

**Figure 2:**
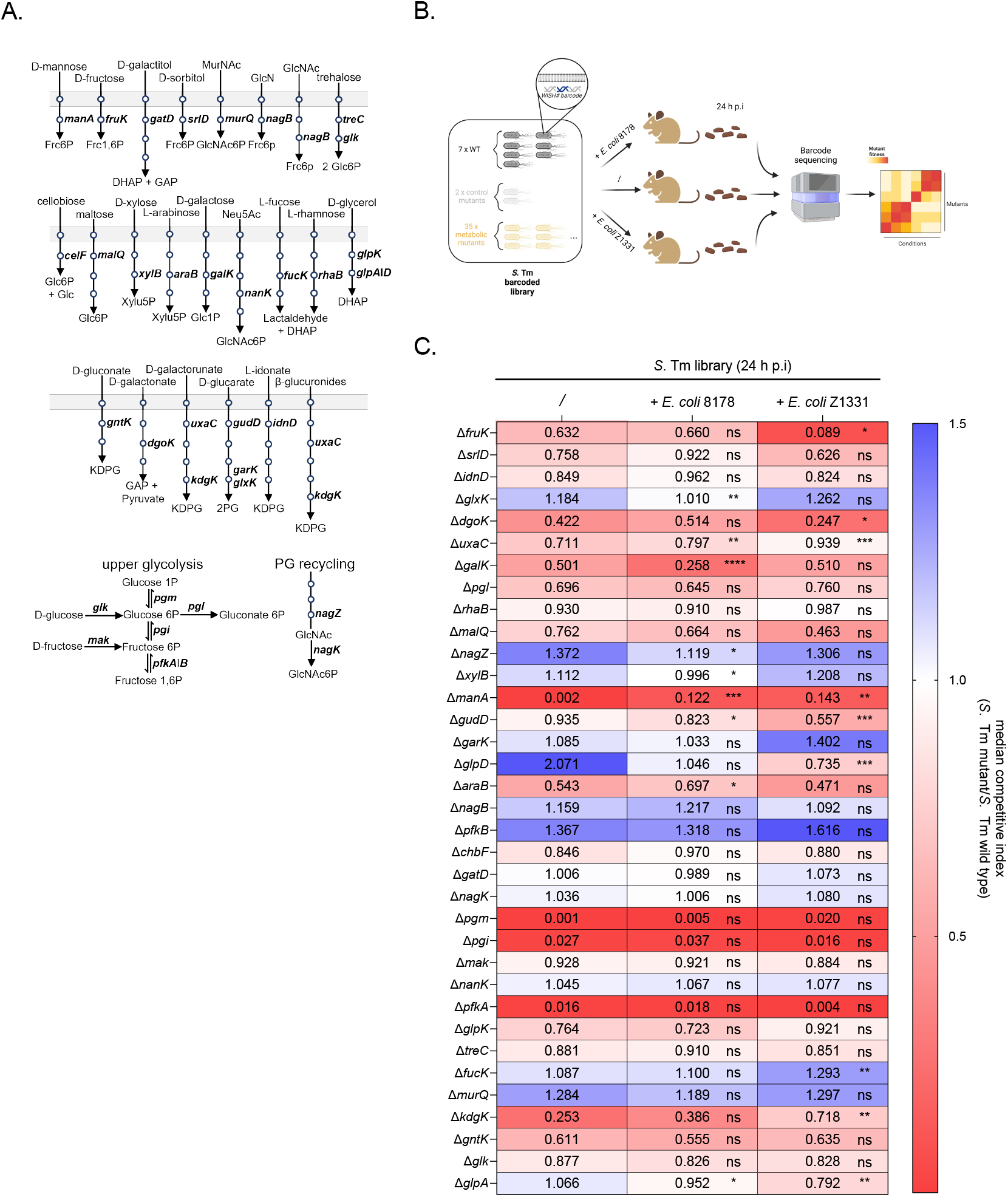
Evaluation of *S*. Tm metabolic requirement in presence of *E. coli* strains. (**A**) Schematic representation of the *S*. Tm metabolic mutant pool, illustrating the targeted carbohydrate utilization pathways, adapted from Schubert et al. (2025). The mutants affect key enzymatic steps, including dehydratases, kinases, and isomerases, positioned between transport reactions and central metabolic intermediates in glycolysis, the Entner-Doudoroff pathway, and the pentose phosphate pathway. Each circle represents an enzymatic step. Abbreviations are listed in Supplementary Table 4 PG, peptidoglycan. (**B**) Experimental scheme. A WISH-barcoded *S*. Tm mutant pool library comprising 7 SL1344 wild type (WT), 2 control mutants (Δ*frd* and Δ*dcuABC*) and 35 deficient strains in carbohydrate utilization was pooled and used in this study. Streptomycin-pretreated SPF 129S6/SvEvTac mice were infected with the *S*. Tm pool alone or in combination with an *E. coli* strain (8178 or Z1331). The fitness of each *S*. Tm mutant was assessed by WISH-barcode sequencing from faecal samples. Created in BioRender. Cherrak, Y. (2025) https://BioRender.com/c18e944. (**C**) Effect of *E. coli* strains on *S*. Tm carbohydrate mutant fitness *in vivo*. The competitive index for each *S*. Tm mutant (listed on the left) is calculated relative to the WT and depicted as a heat map, across 3 different conditions: without *E. coli* (/), in presence of *E. coli* 8178 (+ *E. coli* 8178) in presence of *E. coli* Z1331 (+ *E. coli* Z1331). Median values from a minimum of 6 mice (n ≥ 6) are shown. Two-tailed Mann– Whitney U tests to compare 2 groups (*S*. Tm library alone vs *S*. Tm library + *E. coli* 8178 or *S*. Tm library alone vs *S*. Tm library + *E. coli* Z1331). *p* ≥ 0.05 not significant (ns), *p* < 0.05 (*), *p* < 0.01 (**), *p* < 0.005 (***), *p* < 0.001 (****).

Consistent with earlier findings, we found that in absence of competing strains, *S*. Tm relies on multiple carbohydrates to efficiently grow in the murine gut (Schubert et al., 2025) (Figure 2C, left column). Specifically, the Δ*pgm*, Δ*pgI*, and Δ*pfkA* mutants, responsible for catalysing the sequential conversion of D-glucose 1-phosphate to D-glucose 6-phosphate (*pgm*), then to D-fructose 6-phosphate (*pgi*), and finally to D-fructose 1,6-bisphosphate (*pfkA*)—exhibited a fitness disadvantage compared to the WT control. This was reflected in competitive index (C.I) values below 1, underscoring the essential role of glycolysis under these conditions. Additionally, mutants deficient in metabolizing arabinose (Δ*araB*), galactose (Δ*galK*), fructose (Δ*fruK*), and mannose (Δ*manA*) also displayed reduced fitness (Figure 2C). This suggests that *S*. Tm primarily depends on these metabolites and targeting one of these carbohydrates could potentially impede early gut establishment of the pathogen. Interestingly, the presence of additional *E. coli* competitor strains significantly alters the fitness of *S*. Tm carbohydrate mutants *in vivo* (Figure 2C). This was particularly true for the fitness of the *S*. Tm glycerol (Δ*glpD*) and mannose (Δ*manA*) metabolic mutants which were affected by both *E. coli* 8178 and *E. coli* Z1331. While these effects were consistent across both *E. coli* strains, we also observed strain-specific changes. Particularly, the fitness of the *S*. Tm Δ*fruK* mutant, while relatively unchanged by *E. coli* 8178 (C.I from 0.632 to 0.588), decreased significantly in the presence of *E. coli* Z1331 (C.I = 0.117). More interestingly, the addition of *E. coli* 8178 significantly impaired the fitness of the *S*. Tm *ΔgalK* mutant, as evidenced by an almost 50% reduction in the competitive index, from 0.501 to 0.261—an effect not observed in the presence of *E. coli* Z1331 (Figure 2C). Given the competitive interaction between *S*. Tm and *E. coli* 8178 (but not *E. coli* Z1331, Figures 1B, C), we decided to focus on *S*. Tm metabolic genes that were exclusively affected by *E. coli* 8178 while remaining unaffected by *E. coli* Z1331. This highlighted *galK*, and broadly, galactose metabolism as a strong candidate governing *E. coli* 8178-mediated competition against *S*. Tm, which we selected for further investigation.

### *E. coli* 8178’s restrictions of *S*. Tm early establishment is centred on galactose

Galactose metabolism is widely conserved across *Enterobacteriaceae* and is considered a core component of their metabolic repertoire (Schubert et al., 2025; Näpflin et al., 2025). This is particularly evident in *E. coli* 8178 and *E. coli* Z1331 strains which we tested and confirmed were capable of using galactose as a carbon source *in vitro* (Supplementary Figure 3A). However, it remains unclear whether galactose consumption enhances intestinal colonization of these strains. To evaluate the role of galactose metabolism in *E. coli* within an *in vivo* context, we generated Δ*galK* mutants in both *E. coli* 8178 and *E. coli* Z1331 backgrounds and assessed their fitness in streptomycin-pretreated 129S6/SvEvTac mice. When inoculated with a 1:1 mixture of WT and *galK*-deficient *E. coli* 8178, the WT strain demonstrated a growth advantage over the Δ*galK* mutant (C.I < 1), indicating that galactose metabolism is important for *E. coli* 8178 colonization of the gut (Figure 3A). In contrast, *E. coli* Z1331 did not show a similar dependency on galactose for intestinal colonization, despite both strains metabolizing galactose *in vitro*. We next explored how the galactose utilization of these *E. coli* strains changes in the presence of *S*. Tm. Under these conditions, the *E. coli* 8178 Δ*galK* mutant showed an even greater fitness disadvantage compared to the condition without *S*. Tm, underscoring the importance of galactose for *E. coli* 8178 growth, especially when *S*. Tm is present (Figure 3A). In contrast, the fitness of the *E. coli* Z1331 Δ*galK* mutant remained neutral and unchanged in the presence of *S*. Tm. To further confirm the capacity of *E. coli* 8178 to actively utilize galactose, we measured and compared *S*. Tm galactose-associated fitness in the presence of either the WT or *galK*-deficient *E. coli* 8178 strain. We hypothesized that co-inoculation with an *E. coli* strain incapable of using galactose *in vivo* would deplete other carbohydrates (besides galactose) even more efficiently that the WT *E. coli* 8178 and thereby accentuate the reliance of *S*. Tm on galactose. This in turn, should further exacerbate the competitive attenuation of the *S*. Tm *galK* mutant compared to WT *S*. Tm. Consistent with this hypothesis, *S*. Tm exhibited a significantly greater dependence on galactose when co-inoculated with a *galK*-deficient *E. coli* 8178 strain. This was shown by a *S*. Tm *galK* competitive index of approximately 0.01, which is 26 times lower than in the presence of the WT *E. coli* 8178 strain (C.I = 0.26) (Figure 3B, Supplementary Figure 3B). Together, these findings demonstrate that galactose metabolism is essential for the gut luminal growth of *E. coli* 8178, suggesting that it may be selectively exploited by this strain during intestinal colonization.

**Figure 3:**
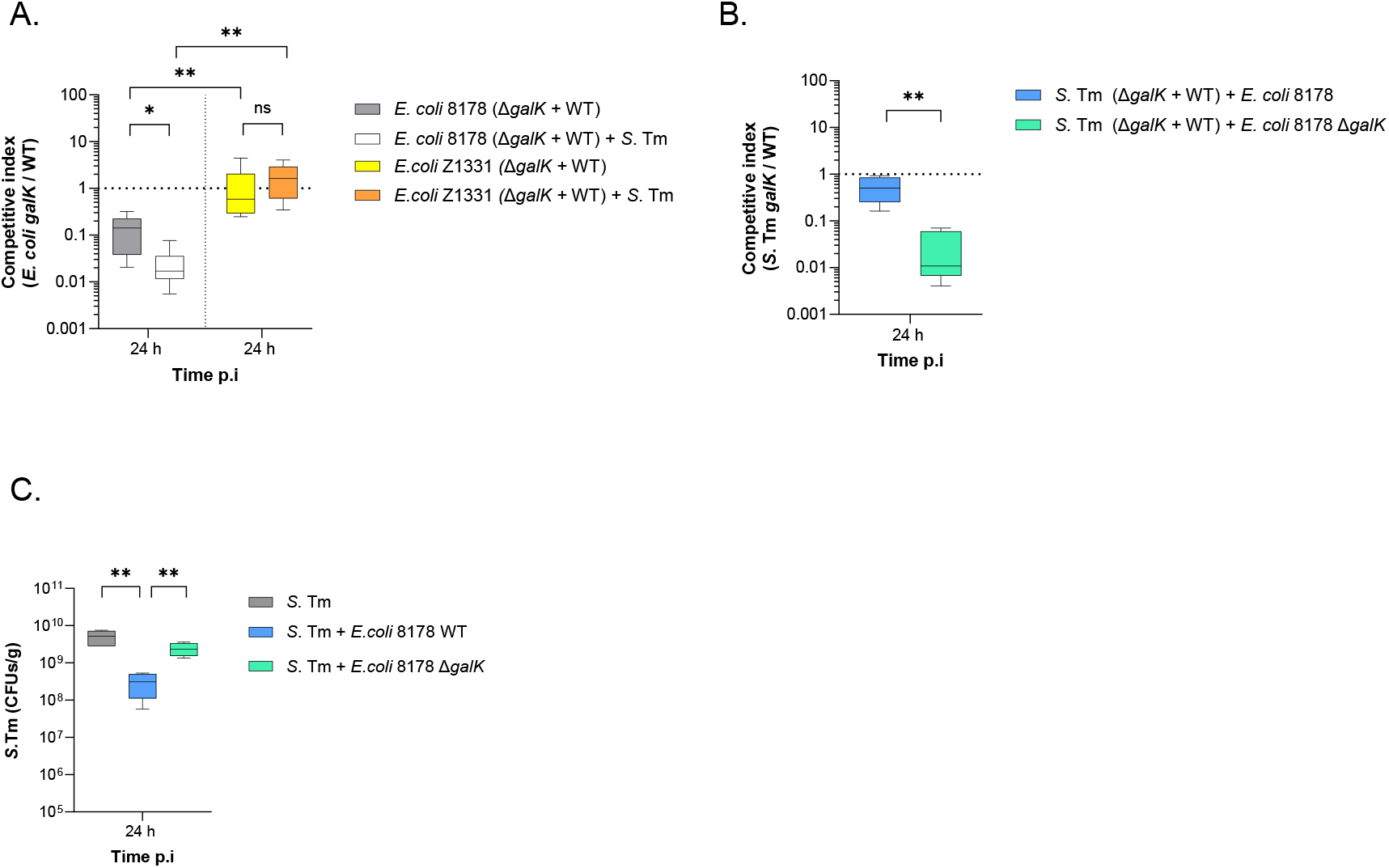
Galactose utilization by *E. coli* 8178 is strain-specific and facilitates the early inhibition of *S*. Tm growth in the gut. **(A)** Fitness of the *galK*-deficient *E. coli* strain *in vivo*. Streptomycin-pretreated 129S6/SvEvTac mice were infected with both the *E. coli* WT and Δ*galK* strains, either in the presence or absence of the WT *S*. Tm (+ *S*. Tm). The competitive index of the Δ*galK* mutant in *E. coli* 8178 (left) and Z1331 (right) at 24 h post-infection (p.i.) are plotted. (**B**) Effect of *E. coli* 8178 on the fitness of *galK*-deficient *S*. Tm *in vivo*. Streptomycin-pretreated 129S6/SvEvTac mice were infected with both the *S*. Tm WT and Δ*galK* strains, in the presence of either the WT or *galK*-deficient *E. coli* 8178. The competitive indexes at 24 h p.i. of the *S*. Tm Δ*galK* mutant are plotted across these two conditions. (**C**) Competitiveness of the *E. coli* 8178 Δ*galK* mutant against *S*. Tm. The *S*. Tm load at 24 h p.i. is plotted and compared between *S*. Tm mono-infected and *S*. Tm + *E. coli* co-infected mice. The strains tested are indicated. (**A-B**) Dotted line: Fitness expected for a fitness-neutral mutation. (**A-C**) All experiments are presented in a box-and-whiskers plot, showing the minimum to maximum values, median, and interquartile range (25th to 75th percentiles). The bar plots show the median. Two-tailed Mann–Whitney U tests to compare 2 groups in each panel. *p* ≥ 0.05 not significant (ns), *p* < 0.05 (*), *p* < 0.01 (**). A minimum of 5 mice (n ≥ 5) were used for each experimental group. CFU: colony forming units.

Based on these findings, we hypothesized that *E. coli* 8178 exploits galactose and consequently attenuates the early gut establishment of *S*. Tm. To test this, we assessed the ability of the *E. coli* 8178 Δ*galK* mutant to restrict *S*. Tm growth in streptomycin-pretreated 129S6/SvEvTac mice. In contrast to the *E. coli* 8178 WT, *S*. Tm colonization was partially restored in mice co-infected with the *E. coli* 8178 *galK* mutant (Figure 3C, Supplementary Figures 3C). This difference was not due to variations in colonization efficacy, as both the *E. coli* 8178 WT and *galK* mutant colonize the gut to a similar extent when competing against *S*. Tm (Supplementary Figure 3D). While our complementation attempts were unsuccessful, this phenotype remained consistent across multiple *galK*-deficient *E. coli* 8178 clones. We also employed a whole genome sequencing approach that ruled out the presence of background mutations in genes other than the *galK* locus. Combined, this indicates that utilization of galactose by *E. coli* 8178 limits *S*. Tm establishment through competitive nutrient exploitation.

## Discussion

Here we report that galactose is a key metabolite in mitigating early gut establishment of *S*. Tm by the mouse commensal *E. coli* 8178. This observation is strain-dependent, as a different *E. coli* isolate, Z1331, did not exhibit this effect.

Characterization of the *S*. Tm growth mitigation was achieved using a previously published, quality-controlled, and rationally designed WISH-barcoded *S*. Tm mutant pool representing the metabolic capacity of *Salmonella* (Schubert et al., 2025). This allowed to highlight how the presence of an additional strain affects *S*. Tm metabolic requirements during gut-luminal growth. Simultaneous testing of multiple mutants within a single animal can efficiently identify important metabolic pathways, reducing costs and the number of animals needed for a comprehensive analysis (Cain et al., 2020). There are two main methodologies for assessing mutant fitness in high-throughput scenarios: randomly barcoded transposon insertion sequencing (RB-TnSeq, van Opijnen et al., 2009) and rationally designed libraries, as used in our study. The latter has technical advantages, utilizing smaller strain libraries for robust mutant representation in niches like the mouse gut (which is often characterized by *S*. Tm population bottlenecks) and enabling simpler bioinformatics, as each mutant is WISH-barcoded for specific gene inactivation. Testing the *S*. Tm mutant pool in competition with different *E. coli* isolates allowed us to report the strain-specific exploitation of galactose as a novel strategy employed by commensal *E. coli* to mitigate *S*. Tm’s early gut invasion.

Freter’s nutrient niche theory suggests that for a bacterium to successfully colonize a niche, it must be the most efficient at metabolizing at least one specific nutrient within that environment (Freter et al., 1983a; Freter et al., 1983b). In recent years, galactitol has been identified as an important substrate for *Enterobacteriaceae*, specifically for *Klebsiella* spp., *E. coli* spp., and *S*. Tm (Oliveira et al., 2020; Eberl et al., 2021; Gül et al., 2023; Osbelt et al., 2024). As a result, galactitol was shown to drive metabolic competition between *S*. Tm and other *Enterobacteriaceae* strains (Eberl et al., 2021; Osbelt et al., 2024). In the current study, we identified galactose as an additional carbohydrate that mediates colonization resistance against *S*. Tm SL1344. Galactitol is a sugar alcohol and the reduction product of galactose. This circumstance requires bacterial oxidation of galactitol before it can be degraded via glycolysis, increasing the metabolic cost of galactitol utilization compared to galactose. Unlike galactitol, which is less common, galactose is relatively abundant in the diet (Englyst et al., 2007; Castillo et al., 2022; Larke et al., 2023) and in the cecum contents of mice with a complex microbiota and gnotobiotic models (Schubert et al, 2025; Nguyen et al, 2024). This is also relevant for the caecal contents of streptomycin-pretreated mice, such as those used in the present study, which were shown to exhibit approximately 2 mM galactose (Nguyen et al., 2024). Taking this information into consideration, we propose that galactose is a key nutrient that can mediate *Enterobacteriaceae*-*Enterobacteriaceae* competition in the gut. In line with this, galactose was shown to act as an important nutrient that promotes *S*. Tm growth in the gut (Stecher et al., 2008). Furthermore, galactose degradation genes in *Enterobacteriaceae* have been identified as part of the core genome, underscoring the significance of galactose as a specific niche for *Enterobacteriaceae* (Schubert et al., 2025). While these observations support the idea of a galactose-rich diet as mediator for *S*. Tm elimination by commensal *Enterobacteriaceae*, it is important to note that galactose supplementation experiments have only partially, but not fully, rescued *S*. Tm survival against *E. coli* 8178. This might be attributable to the host’s relatively high efficiency in absorbing galactose or to the already sufficient levels of galactose in the caecal contents of our mice, making galactose supplementation ineffective. Despite both *E. coli* 8178 and *E. coli* Z1331 having similar *in vitro* growth on galactose as the sole carbon source, only *E. coli* 8178 was able to exploit it and attenuate *S*. Tm growth in the gut. This suggests that the metabolic repertoire of a given bacterial strain alone is insufficient to predict colonization or competitive interactions *in vivo*. The challenge of defining bacterial nutrient preferences *in vivo* stems not only from our limited understanding of the bacterial factors that drive the hierarchical consumption of carbon substrates (Okano et al., 2020) but also from the highly complex and dynamic environment that bacteria encounter within the gut microbiota (Zeng et al., 2022). Although both *E. coli* 8178 and Z1331 can metabolize galactose under defined *in vitro* conditions—where D-galactose is the sole and abundant carbon source—translating this into the gastrointestinal setting may reveal differential requirements for galactose metabolism and other growth-fuelling metabolites. Metabolic exploitation can also be influenced by the sequence of the transcriptional regulator which may differ at the strain level. In some *E. coli* serovars, the start codon of the transcriptional regulator LacI for lactose metabolism is GTG, leading to lower expression of the repressor. This results in higher basal expression of the lactose-utilizing genes, giving these strains a metabolic advantage over isogenic strains with the ATG start codon in the *lacI* gene (Cherrak et al., 2024b). A similar observation was made in *Clostridioides difficile*, in which the transcriptional regulator for trehalose in some epidemic ribotypes has acquired a mutation that allows them to metabolize trehalose at lower concentrations (Collins et al., 2018). Both adaptations provide a competitive advantage in colonization and highlight the extent of strain adaptation required to efficiently utilize a specific niche. Understanding nutrient preferences at the strain-specific level could introduce novel concepts in basic microbiology while also providing optimized strategies for enhancing probiotic efficiency. Taken together, our data indicate that galactose is a crucial nutrient niche for the establishment of *S*. Tm and its exploitation can be affected by the presence of endogenous and specific *E. coli* strains. Similar observations will guide the design of optimized microbiota-based strategies aimed at limiting these nutrients to inhibit or even prevent pathogen invasion.

## Acknowledgment

We would like to thank members of the Hardt group for their comments on the paper. We also thank the EPIC RCHCI staff for support of the animal work. This work has been funded by grants from the Swiss National Science Foundation: grant 310030_192567, Nr. 10.001.588 and the National Centre of Competence in Research (NCCR) Microbiomes (51NF40_225148) to W.-D.H. Y.C. is supported by an EMBO long-term fellowship (ALTF-234-2020) and a flexibility grant from the SNF/NCCR Microbiome (51NF40_180575). C.S is supported by a German Research Foundation fellowship (SCHU 3606/1-1). L.L. is supported by a grant (LA 4572/1-1) from the German Research Foundation. Figures 1A and 2B were created with BioRender.com.

## Contribution

C.S., Y.C., and W.-D.H. conceived and designed the experiments. C.S. trained and supervised J.N. C.S and J.N. performed the *in vivo* screens and analysis. Y.C. trained and supervised L.P., and both performed the *S*. Tm – *E. coli* competitive experiments. L.L generated the *galK* mutant in the *E. coli* Z1331 background. Y.C. and C.S. wrote the manuscript with the contributions from all authors.

## Materials availability

Mouse lines used in this study can be obtained from Jackson laboratories.

## Data availability

All data needed to evaluate the conclusions of this study are presented in the Article and Supplementary Information. The amplicon sequencing data for the WISH-barcoded S. Typhimurium mutant pool experiments has been deposited in the European Nucleotide Archive under the accession number: PRJEB85163.

## Methods

### Animals and ethic statements

Male and female 8 to 12 weeks old 129S6/SvEvTac (Jackson Laboratory) mice were randomly assigned to experimental groups and used in this study. The mice were housed under SPF conditions in individually ventilated cages at the EPIC mouse facility, ETH Zurich. All animal experiments were conducted in accordance with Swiss and cantonal regulations and were reviewed and approved by the Tierversuchskommission, Kantonales Veterinäramt Zürich under license ZH158/2019, ZH108/2022, and ZH109/2022, in compliance with the cantonal and Swiss legislation.

### Strains, media, and chemicals

All strains, plasmids and oligonucleotides used in this study are listed in Tables S1-S3. Custom oligonucleotides were synthetized by Microsynth AG (Balgach, Switzerland). *E. coli* 8178 and Z1331 strains originate from previous work (Stecher *et al*., 2012; Wotzka *et al*., 2018). The *S*. Tm mutant pool was designed in Schubert et al., 2025. Bacterial strains were routinely cultured in lysogeny broth (LB) with or without 1% agar-agar. Mutant selection was carried out using antibiotics: streptomycin (100 μg/ml), carbenicillin (100 μg/ml), kanamycin (50 μg/ml), and chloramphenicol (30 μg/ml). Gene deletions in *E. coli* 8178 and Z1331 were performed using a modified lambda red recombinase one-step inactivation method (Datsenko & Wanner, 2000). In brief, kanamycin resistance cassettes (kanamycin for deletions, were PCR-amplified (*Phusion* DNA Polymerase, Sigma-Aldrich)) with primers containing 50-nucleotide homologous regions flanking the target site. Electroporation of these PCR products into *E. coli* cells expressing lambda red recombinase from the pSIM5 plasmid allowed for mutant generation, which were then selected on antibiotic plates (Dinner et al., 2011). Gene deletions were confirmed by colony PCR and whole-genome sequencing, with a particular focus on the *E. coli* 8178 Δ*galK* mutant (PRJEB85163).

### Mouse infection experiments

The 8-to 12-week-old mice were orally pretreated with streptomycin (25 mg) 24 h before inoculation (Barthel et al., 2003). *S*. Tm and *E. coli* cultures were individually grown in LB at 37°C for 4 h and washed twice with a phosphate-buffered saline solution (PBS: 137 mM NaCl, 2.7 mM KCl, 10 mM Na_2_HPO_4,_ and 1.8 mM KH_2_PO). The *S*. Tm pool was cultured in a similar fashion from pre-mixed cryo stocks. Prior to colonization experiments, *E. coli* strains were electroporated with the pRSF1010 (P3) plasmid from *Salmonella enterica* serovar Typhimurium SL1344 which confers streptomycin resistance (Guerry et al., 1974). Each mouse received a single oral dose of 50 μl containing approximately 5 × 10^7^ colony forming units (cfu) of an inoculum mixture consisting of equal ratios of the indicated strains. Faecal samples were collected 24 hours post-infection, and animals were euthanized via CO_2_ asphyxiation on day 1 post-infection. Faecal samples were then suspended in 1 ml of PBS and homogenized using a TissueLyser (Qiagen). Bacterial loads were quantified by plating the homogenized suspension on selective MacConkey or LB agar plates. In competition assays involving two bacterial strains (WT and mutant), the mutant’s bacterial load was directly quantified by plating on antibiotic-containing agar. Conversely, the bacterial load of the WT strain was derived from the total colony count on antibiotic-free plates using the formula: cfu_WT_ = cfu_total_ − cfu_mutant_. The load of each mutant strain was then normalized to its initial inoculum, and this value was used to calculate the normalized competitive index (C.I.) to assess its fitness relative to the WT strain. The C.I. for each mutant was determined by taking the ratio of cfu_mutant_ to cfu_WT_, divided by the ratio of the mutant and WT strains in the inoculum.

### Evaluation of *S*. Tm metabolic mutant fitness *via* barcode counting

Preparation and analysis of *in vivo* samples containing the *S*. Tm WISH barcoded mutant pool was achieved as previously reported (Schubert et al., 2025). Specifically, faecal *S*. Tm cells were cultivated in LB medium supplemented with carbenicillin to select for WISH-barcoded strains. Following enrichment, cells were pelleted and stored at -20°C. DNA was extracted from the thawed pellets using the Qiagen Mini DNA kit, following the manufacturer’s protocol. Amplification of WISH barcodes *via* PCR was performed using the DreamTaq MasterMix (Thermo Fisher Scientific) under the following cycling conditions: (1) initial denaturation at 95°C for 3 min, (2) denaturation at 95°C for 30 sec, (3) annealing at 55°C for 30 sec, (4) extension at 72°C for 20 sec, repeated for 25 cycles, followed by (5) a final extension at 72°C for 10 min. The resulting PCR products were purified (Macherey-Nagel) and subsequently indexed by a second PCR with unique dual index primers using the following program: (1) 95°C for 3 min, (2) 95°C for 30 sec, (3) 55°C for 30 sec, (4) 72°C for 20 sec, repeated for 10 cycles, with a (5) final extension at 72°C for 3 min. The indexed products were checked by gel electrophoresis (1% w/v agarose in Tris, acetate, EDTA buffer: 40 mM tris-acetate, 1 mM EDTA), pooled based on band intensity, and purified using AMPure beads (Beckman Coulter) for library preparation. Amplicon sequencing was carried out by BMKGENE (Münster, Germany) using the Novaseq platform, with 150 bp paired-end reads at a target output of 1 G per sample. Following sequencing, reads were demultiplexed and WISH-barcodes were counted using the mBARq software (Sintsova et al., 2024). Misreads or mutations of up to five bases were assigned to the closest matching WISH-tag sequence. WISH-barcode counts for each mouse are provided in Source data file. These counts were then used to calculate competitive fitness and Shannon evenness scores across seven wild-type strains. WISH counts ≤ 10 were excluded from further analysis, setting the detection limit as previously described by Daniel et al. (2024).

### *In vitro* galactose utilization assay

Biolog PM1 MicroPlates (BIOLOG; Bochner et al., 2001) were used for metabolic profiling of *S*. Tm and *E. coli* strains according to the manufacturer’s protocol. Bacteria were grown overnight in M9 minimal medium with 25 mM pyruvate as the sole carbon source at 37°C with shaking. They were then washed and resuspended in M9 medium without pyruvate to an OD_600_ of 0.056. 100 μl of this suspension was added to each well of the microplate. Plates were sealed with parafilm and incubated at 37°C. OD_600_ was measured every 10 minutes for 12 hours with orbital shaking at 282 rpm using a BioTek Synergy H1 Microplate spectrophotometer (Agilent).

### Statistical analysis

Statistical analyses and data visualization were conducted using GraphPad Prism version 9.2.0 for Windows (GraphPad Software, La Jolla, CA, USA). For comparisons between two groups, the unpaired Mann–Whitney U-test was employed to evaluate statistical significance, based on rank comparison. *P* values < 0.05 were considered to indicate statistical significance.

## Supplementary Figures captions

**Supplementary Figure 1:**
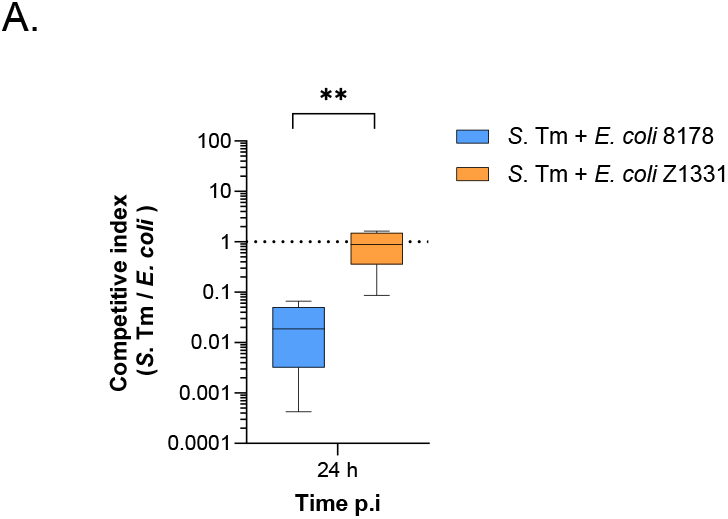
*E. coli* 8178 and Z1331 loads in co-infected mice. (**A**). The competitive index between *S*. Tm and *E. coli* from Figures 1B, C are shown. The box-and-whiskers plot represent the minimum to maximum values, median, and interquartile range (25th to 75th percentiles). Two-tailed Mann–Whitney U tests to compare 2 groups in each panel. *p* < 0.01 (**).

**Supplementary Figure 2:**
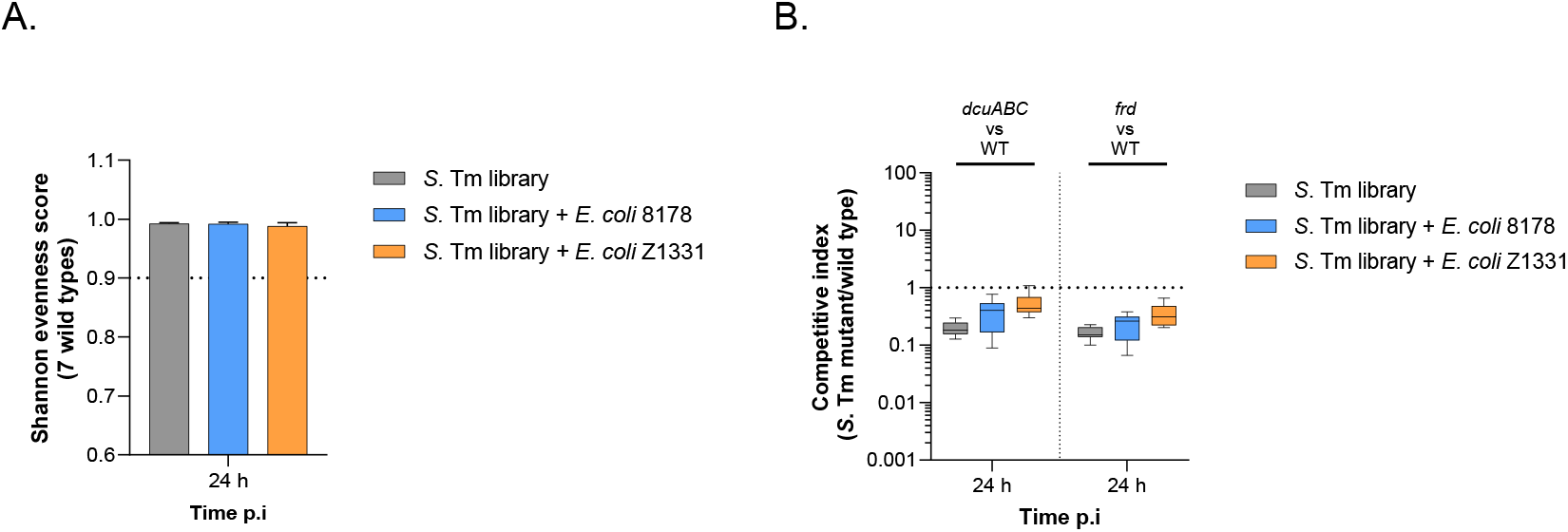
Quality-control of the *S*. Tm mutant pool analysis *in vivo*. (**A**) The Shannon evenness score at 24 h post-infection (p.i) is presented for each condition tested. A score close to 1 signifies an even distribution of the 7 *S*. Tm wild-type strains, reflecting the absence of a bottleneck effect. The median and standard deviation are shown, with the dotted line representing the threshold (0.9) below which samples were excluded from analysis. (**B**) Fitness of the internal *S*. Tm metabolic mutant controls. The competitive index of the *S*. Tm Δ*dcuABC* (left) and Δ*frd* (mutant) at 24 h p.i is plotted for each condition tested. Data are presented in a box-and-whiskers plot, showing the minimum to maximum values, median, and interquartile range (25th to 75th percentiles). The bar plots show the median. Dotted line: competitive index expected for a fitness-neutral mutation. (**A-B**) A minimum of 6 mice (n ≥ 6) were used for each condition tested.

**Supplementary Figure 3:**
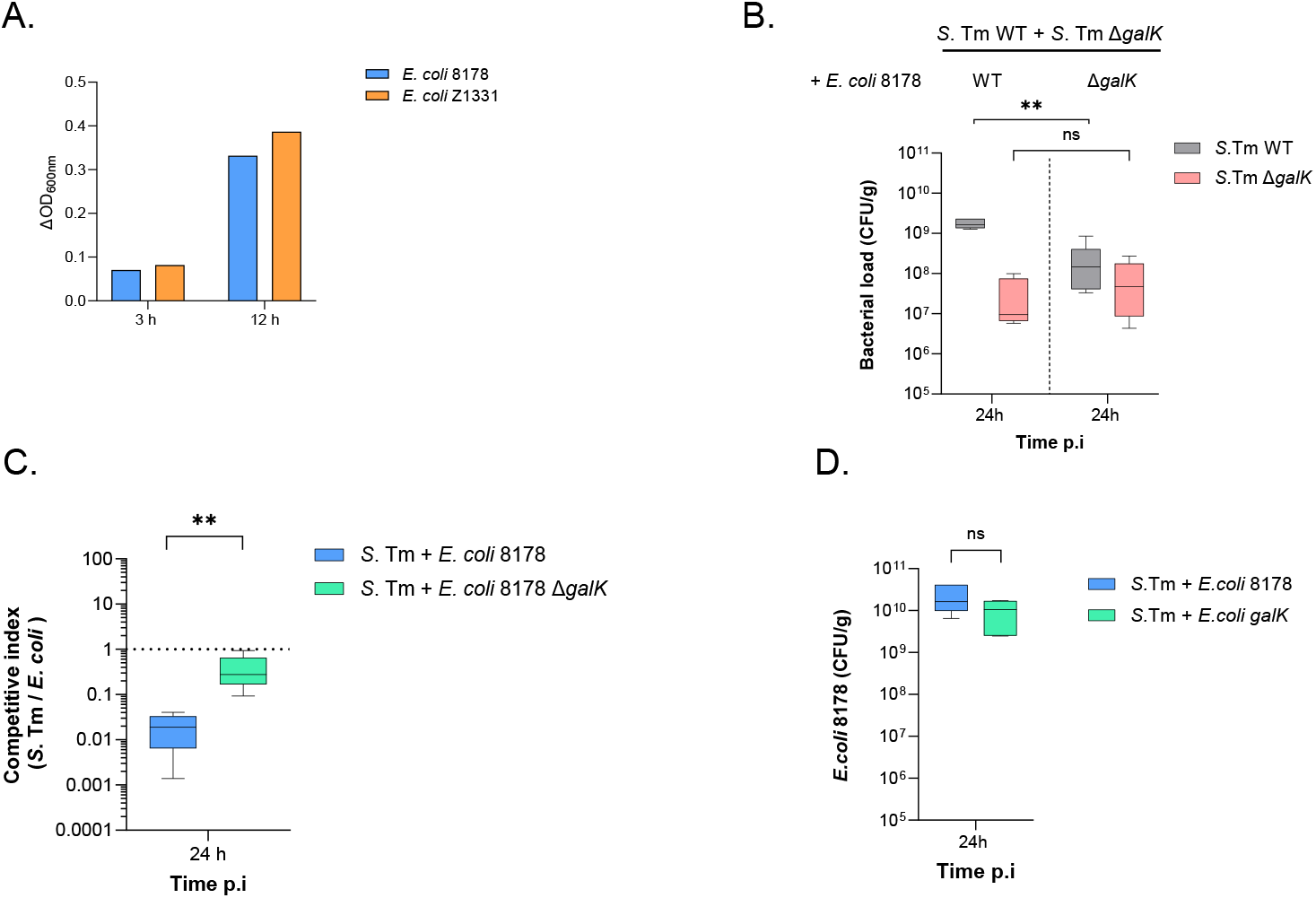
*E. coli* 8178 and Z1331 grow on galactose as the sole carbon source. (**A)** *In vitro* growth on galactose as the sole carbon source was assessed for *E. coli* 8178 and Z1331 strains. Growth was measured in a Biolog PM1 microplate, with data plotted as ΔOD_600nm_ for 3 hours and 12 hours for each bacterium as indicated. (**B**) Bacterial loads of the WT and *galK*-deficient *S*. Tm strains in presence of the WT (left) or Δ*galK* (right) *E. coli* 8178 are plotted as CFU per gram of feces at 24 h p.i (from Figure 3B). (**C**) Competitive index at 24 h p.i between *S*. Tm and *E. coli* (from Figure 3C). **(D**) *E. coli* 8178 loads in *S*. Tm + *E. coli* co-infected mice (from Figure 3C). (**B-D**) Data are presented in a box-and-whiskers plot, showing the minimum to maximum values, median, and interquartile range (25th to 75th percentiles). The bar plots show the median. Two-tailed Mann–Whitney U tests to compare 2 groups in each panel. *p* ≥ 0.05 not significant (ns), *p* < 0.01 (**).

## Supplementary Tables

**Table S1: Bacterial strains used in this study**

**Table S2: Plasmid used in this study**

**Table S3: Oligonucleotides used in this study**

